# Modeling COVID-19 disease biology to identify drug treatment candidates

**DOI:** 10.1101/2022.04.18.488660

**Authors:** Walter J. Jessen, Stefan Diaz Gaisenband, M’Lissa Quintanilla, Sadiq Lula, Patrick McLeroth, Adam Sullivan, Stanley Letovsky

## Abstract

Coronavirus disease 2019 (COVID-19) is an infectious disease caused by severe acute respiratory syndrome coronavirus 2 (SARS-CoV-2). Currently, there are a limited number of effective treatments. A variety of drugs that have been approved for other diseases are being tested for the treatment of COVID-19, and thus far only remdesevir, dexamethasone, baricitinib, tofacitinib, tocilizumab, and sarilumab have been recommended by the National Institutes of Health (NIH) COVID-19 Treatment Guidelines Panel for the therapeutic management of hospitalized adults with COVID-19. Using a disease biology modeling approach, we constructed a protein-protein interactome network based on COVID-19- associated genes/proteins described in research literature together with known protein-protein interactions in epithelial cells. Phenotype and disease enrichment analysis of the COVID-19 disease biology model demonstrated strong statistical enrichments consistent with patients’ clinical presentation. The model was used to interrogate host biological response induced by SARS-CoV-2 and identify COVID-19 drug treatment candidates that may inform on drugs currently being evaluated or provide insight into possible targets for potential new therapeutic agents. We focused on cancer drugs as they are often used to control inflammation, inhibit cell division, and modulate the host microenvironment to control the disease. From the top 30 COVID-19 drug candidates, twelve have a role as an antineoplastic agent, seven of which are approved for human use. Altogether, nearly 40% of the drugs identified by our model have been identified by others for COVID-19 clinical trials. Disease biology modeling incorporating disease-associated genes/proteins discussed in the research literature together with known molecular interactions in relevant cell types is a useful method to better understand disease biology and identify potentially effective therapeutic interventions.

## Introduction

As of the 21^st^ of February, 2022, the coronavirus disease 2019 (COVID-19) has caused over 422,188,754 infections and killed over 5,876,766 people worldwide ^1^. The SARS-CoV-2 virus that causes COVID-19 was first identified in Wuhan, China, as a novel coronavirus that was related to a cluster of pneumonia cases most commonly associated with respiratory and gastrointestinal symptoms, but can vary in symptoms based on disease severity ^2^. While COVID-19 primarily affects the respiratory system, the virus can affect any organ in the body; in critically ill patients, multiple organs are often affected ^3^. Most infections will be asymptomatic or mild; however, many people will suffer from moderate to severe infections and a minority will develop acute respiratory distress syndrome (ARDS) requiring critical care support in an intensive care setting.

An epidemiological study across eighteen European countries in 1,420 patients with mild-to-moderate COVID-19 who did not require hospitalization in intensive care units showed that patients with non-severe disease most commonly suffer from headache (70.3%), loss of smell (70.2%), nasal obstruction (67.8%), cough (63.2%), asthenia (63.3%), myalgia (62.5%), rhinorrhea (60.1%), gustatory dysfunction (54.2%), sore throat (52.9%), and fever (45.4%). However, these symptoms were reported to vary significantly depending on age and sex ^4^. Several small studies in early 2020 across local health centers in China revealed that patients admitted to hospital with severe COVID-19 presented most often fever (94-98%), cough (76-79%), dyspnea (55%), and myalgia or fatigue (23-44%) ^5, 6^. During the peak of the pandemic, the number of hospitalized patients increased to unsustainable levels across all countries globally, which led to national healthcare systems being overwhelmed and unable to cope with patient volume. Due to the critical situation, the industry had to urgently address unmet medical needs and turned towards drugs that were readily available in the market for the treatment of other conditions. Several large clinical trials have been conducted with marketed drugs, especially in patients that were admitted to the hospital presenting with greater disease severity, and have provided invaluable additional information on the clinical features of COVID-19 and patients’ baseline characteristics. Examples of drugs previously trialed and currently available either under emergency use authorization (EUA) or full approval for the treatment of hospitalized COVID-19 patients that have been repurposed from their use in other indications are baricitinib, a reversible Janus kinase (JAK) inhibitor for treatment of rheumatoid arthritis in adults whose disease was not well controlled by tumor necrosis factor (TNF) inhibitors; dexamethasone, an anti-inflammatory steroid used for many different inflammatory conditions such as allergic disorders and skin conditions; remdesevir, a broad-spectrum antiviral originally developed to treat hepatitis C; tocilizumab, a humanized monoclonal antibody against the interleukin-6 receptor (IL-6R) for the treatment of rheumatoid arthritis and systemic juvenile idiopathic arthritis; tofacitinib, another JAK inhibitor for treatment of rheumatoid arthritis, psoriatic arthritis and ulcerative colitis; and sarilumab, an injectable prescription medicine called an IL-6R blocker for treatment of adults with moderate to severe active rheumatoid arthritis not responding or tolerating treatment with at least one disease-modifying antirheumatic drug ^7^. In addition to these drugs, many other compounds have been used in a clinical trial setting to investigate whether they could be used as a potential treatment in COVID-19 infected patients; however, only a minority will be authorized for treatment. Given the global health crisis, more sophisticated ways to develop new drugs against COVID-19 are being explored.

We have developed a new methodology to repurpose readily available drugs for a given disease and/or identify candidate biomarkers. This methodology is based on *in silico* disease biology modelling which aims to recapitulate the molecular environment of diseased target cells to identify candidate biomarkers and/or drugs that can be proposed as options for disease treatment. Others studying SARS-CoV-2 infection and COVID-19 have used protein-protein interaction (PPI) networks to analyze host-virus protein-protein interaction ^8–10^, describe the interactome of the coronavirus S-glycoprotein and host proteins ^11^, investigate viral-viral and virus-host PPI networks in peripheral blood mononuclear cells ^12^, analyze the immune system PPI network and review potential therapeutic targets for drug repurposing against COVID-19 ^13^, and identify potentially repurposable drugs to treat COVID-19 ^14–16^.

Here, we describe the development and analysis of a COVID-19 disease biology model based on genes/proteins significantly associated with the disease discussed in the research literature together with known molecular interactions in epithelial cells to better understand host biological response induced by SARS-CoV-2 and identify effective interventions. We review impacted biological processes/pathways and drug candidates that could be repurposed to treat COVID-19 infection.

## Methods

### Model construction

To build an interactome network, we first identified relationships between COVID-19 disease and genes/proteins in Medline and PubMed Central using PolySearch 2.0, an online text-mining system ^17^. COVID-19 associations were mined on April 6th, 2021; influenza A H1N1 associations were mined on June 1st, 2021.To ensure consistent identification, our query for COVID-19 included the manually added synonyms coronavirus disease 2019, covid-19 infection, sars-cov-2 infection, and severe acute respiratory syndrome coronavirus 2, and for influenza A H1N1, our query (influenza A H1N1) included the manually added synonyms A/H1N1, swine flu, and H1N1. Default settings were used for both custom filter words and custom negation words. Following both automated mapping and manual review, we identified 14 COVID-19 and 13 influenza A H1N1 genes/proteins, which were used as seeds to construct interactome networks in Ingenuity Pathway Analysis (IPA) (QIAGEN Inc., https://digitalinsights.qiagen.com/products-overview/discovery-insights-portfolio/analysis-and-visualization/qiagen-ipa/) ^18^.

Interactome networks were generated using established algorithms and settings, identical for both COVID-19 and influenza A H1N1, based primarily on direct physical molecular interactions. We start by growing up- and down-stream of the seeds, and then look for direct physical interactions between found molecules. We then apply the shortest paths algorithm to the largest single multi-node connected subnetwork and look to add one new molecule (e.g. shortest paths +1) that has been identified experimentally to physically interact with any of the nodes and found molecules.

Influenza A H1N1 is known to infect both epithelial and immune cells ^19, 20^. Thus, the only difference in model construction was cell type: COVID-19 used PPI knowledge identified in epithelial cells ^21–23^ while influenza A H1N1 used PPI knowledge identified in either epithelial or immune cells.

### Gene list enrichment analysis

We performed gene list enrichment analysis for the Human Phenotype Ontology (HPO), diseases (DisGeNET BeFree), Gene Ontology Biological Process (GO:BP), Pathway Ontologies (MSigDB C2 BIOCART (v7.3) and PantherDB), and Drugs (Comparative Toxicogenomics Database (CTD) and Stitch) using ToppFun, a module of the ToppGene Suite (https://toppgene.cchmc.org) ^24^. Correction for multiple testing was performed using the Benjamini-Hochberg (BH) procedure ^25^ to calculate a false discovery rate (FDR) adjusted p-value (q-value FDR BH). Human phenotypes, diseases, GO:BPs, pathways and drugs were considered significantly enriched if the q-value for the overlap comparison test between input model genes and a given ontology class was less than 0.05. Enrichment analysis typically identifies multiple terms down the same branch of an ontology directed acyclic graph (DAG); nodes further down the DAG provide more information but are less significant and have fewer associated genes than higher nodes. For the top 500 GO:BP terms, we used REVIGO to find a representative subset and group semantically similar terms together ^26^. We then selected representative terms from each of the large semantically similar clusters. For diseases, we categorized the top 100 DisGeNET BeFree terms using the Disease Ontology (https://disease-ontology.org/) ^27^. For all additional enrichment analyses, to ensure prudent selection of significantly enriched terms and to minimize over-representation, terms were manually reviewed to select those that best balanced context, informative content, and gene enrichment.

### Disease enrichment mapping

To objectively and systematically compare and contrast disease classes instead of individual conditions between COVID-19 and influenza A H1N1 models, the top 100 disease enrichments for each model were mapped from DisGenNET BeFree names to Disease Ontology IDs (DOIDs) ^27^. The parent Disease Ontology class(es) was/were used to represent each condition (second level term in the Disease Ontology tree).

### Clinical trials

Clinical trial information was obtained from Informa Pharma Intelligence’s Trialtrove and Clinicaltrials.gov, accessed January 28^th^, 2022. To search for studies in Trialtrove, the query (Disease is Infectious Disease: Novel coronavirus (2019-nCoV, COVID-19)) AND [(Tested Drug is X)] was used, which included any alternative names for the drug. Search results were then manually sorted to eliminate any non-relevant results. To search for studies in Clinicaltrials.gov, COVID-19 was entered as the ‘condition or disease’ and the drug name was entered under ‘Intervention/treatment’. Search results were then manually sorted to eliminate any non-relevant results. The results from both searches were then combined and de-duplicated for each drug.

## Results

The COVID-19 disease biology model consists of 31 gene/protein nodes (**Figure 1**). After discarding unconnected nodes and small multi-node fragments, of the 14 genes/proteins identified in the literature, six remained in the single large multi-node interactome network. The eight genes/proteins that were discarded included ACE2, SPINK5, FKBP1A, LTB, CEP250, C5AR1, WDHD1, and MIR410; no molecular interactions with these genes/proteins specific to epithelial cells were identified with any other node in the network. Using the same parameters for literature mining and interactome construction, we also built a disease biology model for influenza A H1N1 as a control.

**Figure 1.**
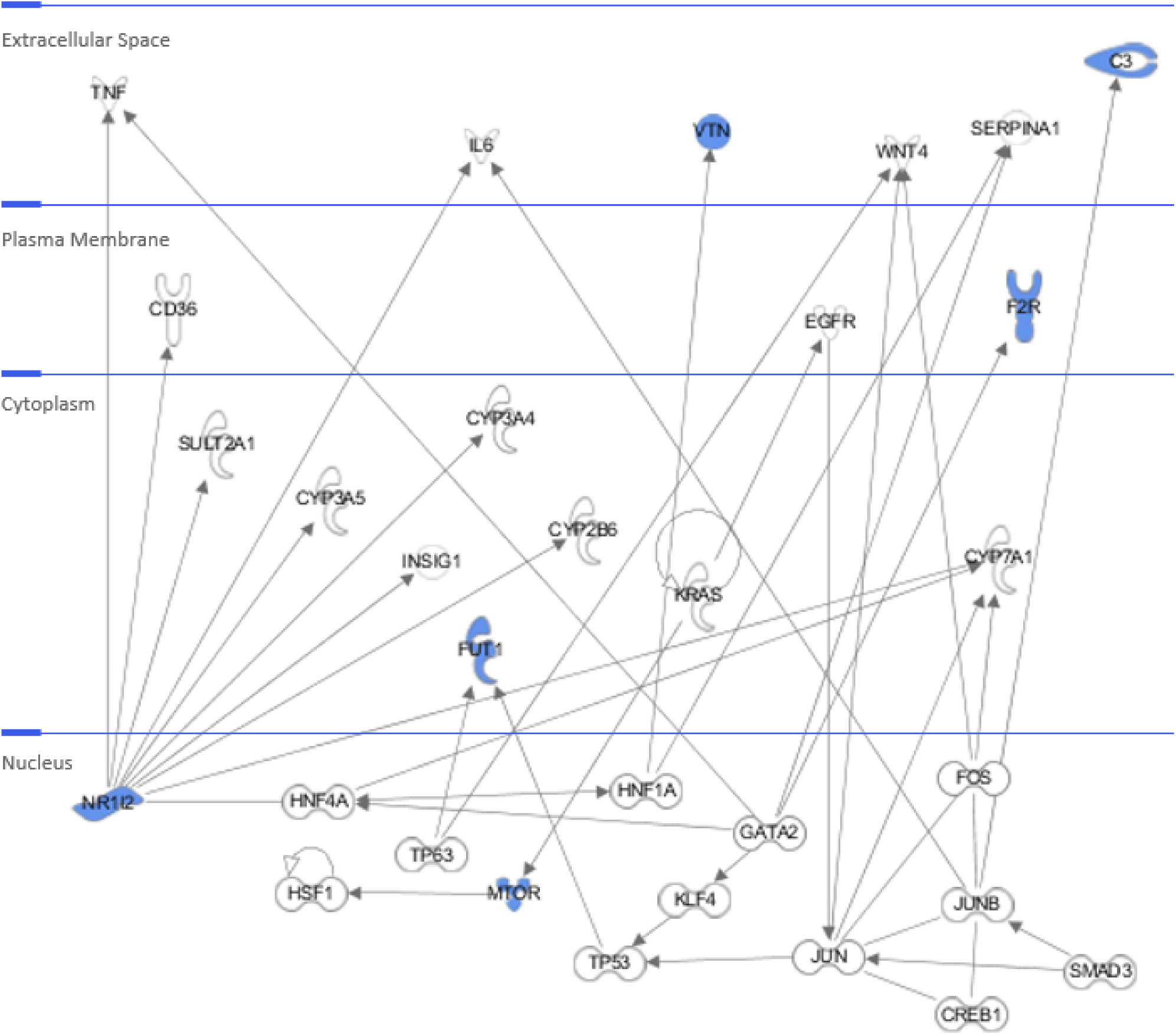
COVID-19 disease biology model. Nodes represent genes/proteins; lines between nodes represent one or more physical molecular interactions identified experimentally. Blue nodes indicate genes/proteins initially identified in the literature that remain in the final model.

### Host biological response induced by SARS-CoV-2

To better understand the biology represented by the COVID-19 model, we performed Gene Ontology (GO) biological process and pathway enrichment analyses. We used REVIGO to remove redundant GO terms for the top 500 significantly enriched biological process terms and grouped semantically similar terms together. For large clusters, we then selected representative terms (**Table 1**). Pathway enrichment analysis incorporated three sources: Gene Map Annotator and Pathway Profiler (GenMAPP), the molecular signatures database (MSigDB C2 BIOCARTA v7.3), and PATHER Pathways (**Table 2**).

**Table 1.** Representative gene ontology biological process terms significantly enriched in the COVID-19 disease biology model. Red gene symbols indicate the COVID-19-associated genes/proteins initially identified in the literature.

| GO:ID | Biological Process Name | q-value<br>FDR B&H | Hit Count in<br>Query List | Hit Count<br>in Genome | Model Genes/Proteins |
| --- | --- | --- | --- | --- | --- |
| GO:0009611 | response to wounding | 8.29E-11 | 15 | 746 | GATA2, JUN, **C3**, SERPINA1, TNF, **VTN**, WNT4, KLF4, EGFR, HNF4A, **F2R**, **MTOR**, IL6, CD36, SMAD3 |
| GO:0006629 | lipid metabolic process | 1.24E-08 | 16 | 1503 | **C3**, HNF1A, CYP2B6, TNF, **NR1I2**, WNT4, EGFR, HNF4A, SULT2A1, CYP3A4, CREB1, CYP3A5, **MTOR**, CYP7A1, CD36, INSIG1 |
| GO:0008202 | steroid metabolic process | 1.61E-08 | 10 | 371 | HNF1A, CYP2B6, TNF, **NR1I2**, WNT4, SULT2A1, CYP3A4, CYP3A5, CYP7A1, INSIG1 |
| GO:0043066 | negative regulation of apoptotic process | 1.61E-08 | 14 | 1058 | GATA2, KRAS, JUN, TNF, WNT4, HSF1, KLF4, EGFR, **F2R**, **MTOR**, IL6, TP63, TP53, SMAD3 |
| GO:0034599 | cellular response to oxidative stress | 8.33E-08 | 9 | 342 | JUN, TNF, HSF1, KLF4, EGFR, FOS, IL6, CD36, TP53 |
| GO:0010817 | regulation of hormone levels | 5.32E-07 | 10 | 618 | HNF1A, TNF, WNT4, EGFR, HNF4A, SULT2A1, CYP3A4, CREB1, CYP3A5, IL6 |
| GO:0002521 | leukocyte differentiation | 5.82E-07 | 10 | 626 | GATA2, JUN, JUNB, TNF, WNT4, CREB1, **MTOR**, FOS, IL6, TP53 |
| GO:0006805 | xenobiotic metabolic process | 7.99E-07 | 6 | 122 | CYP2B6, **NR1I2**, HNF4A, SULT2A1, CYP3A4, CYP3A5 |
| GO:0070372 | regulation of ERK1 and ERK2 cascade | 1.16E-06 | 8 | 356 | JUN, **C3**, TNF, KLF4, EGFR, **F2R**, IL6, CD36 |
| GO:0050817 | coagulation | 1.37E-06 | 8 | 368 | GATA2, **C3**, SERPINA1, **VTN**, HNF4A, **F2R**, IL6, CD36 |
| GO:0070371 | ERK1 and ERK2 cascade | 1.59E-06 | 8 | 377 | JUN, **C3**, TNF, KLF4, EGFR, **F2R**, IL6, CD36 |
| GO:0000165 | MAPK cascade | 3.17E-06 | 11 | 1020 | KRAS, JUN, **C3**, TNF, WNT4, HSF1, KLF4, EGFR, **F2R**, IL6, CD36 |
| GO:0006954 | inflammatory response | 6.39E-06 | 10 | 862 | **C3**, SERPINA1, TNF, KLF4, EGFR, **F2R**, FOS, IL6, CD36, SMAD3 |
| GO:0002682 | regulation of immune system process | 1.09E-05 | 13 | 1747 | GATA2, KRAS, JUN, **C3**, TNF, **VTN**, CREB1, **MTOR**, FOS, IL6, CD36, TP53, SMAD3 |
| GO:0046427 | positive regulation of receptor signaling pathway via JAK-STAT | 1.37E-05 | 4 | 49 | TNF, HNF4A, **F2R**, IL6 |
| GO:0050890 | cognition | 1.81E-05 | 7 | 375 | KRAS, JUN, TNF, EGFR, CREB1, **MTOR**, FOS |
| GO:0006952 | defense response | 2.71E-05 | 13 | 1935 | KRAS, **C3**, SERPINA1, TNF, HSF1, KLF4, EGFR, **F2R**, FOS, IL6, CD36, TP53, SMAD3 |
| GO:2000334 | positive regulation of blood microparticle formation | 2.80E-05 | 2 | 2 | TNF, CD36 |
| GO:1905153 | regulation of membrane invagination | 3.63E-05 | 3 | 19 | GATA2, **C3**, CD36 |
| GO:0001817 | regulation of cytokine production | 5.68E-05 | 9 | 897 | **C3**, TNF, HSF1, KLF4, **F2R**, CREB1, IL6, CD36, SMAD3 |
| GO:0045321 | leukocyte activation | 5.71E-05 | 11 | 1447 | GATA2, JUN, **C3**, SERPINA1, TNF, WNT4, **MTOR**, IL6, CD36, TP53, SMAD3 |

**Table 2.**
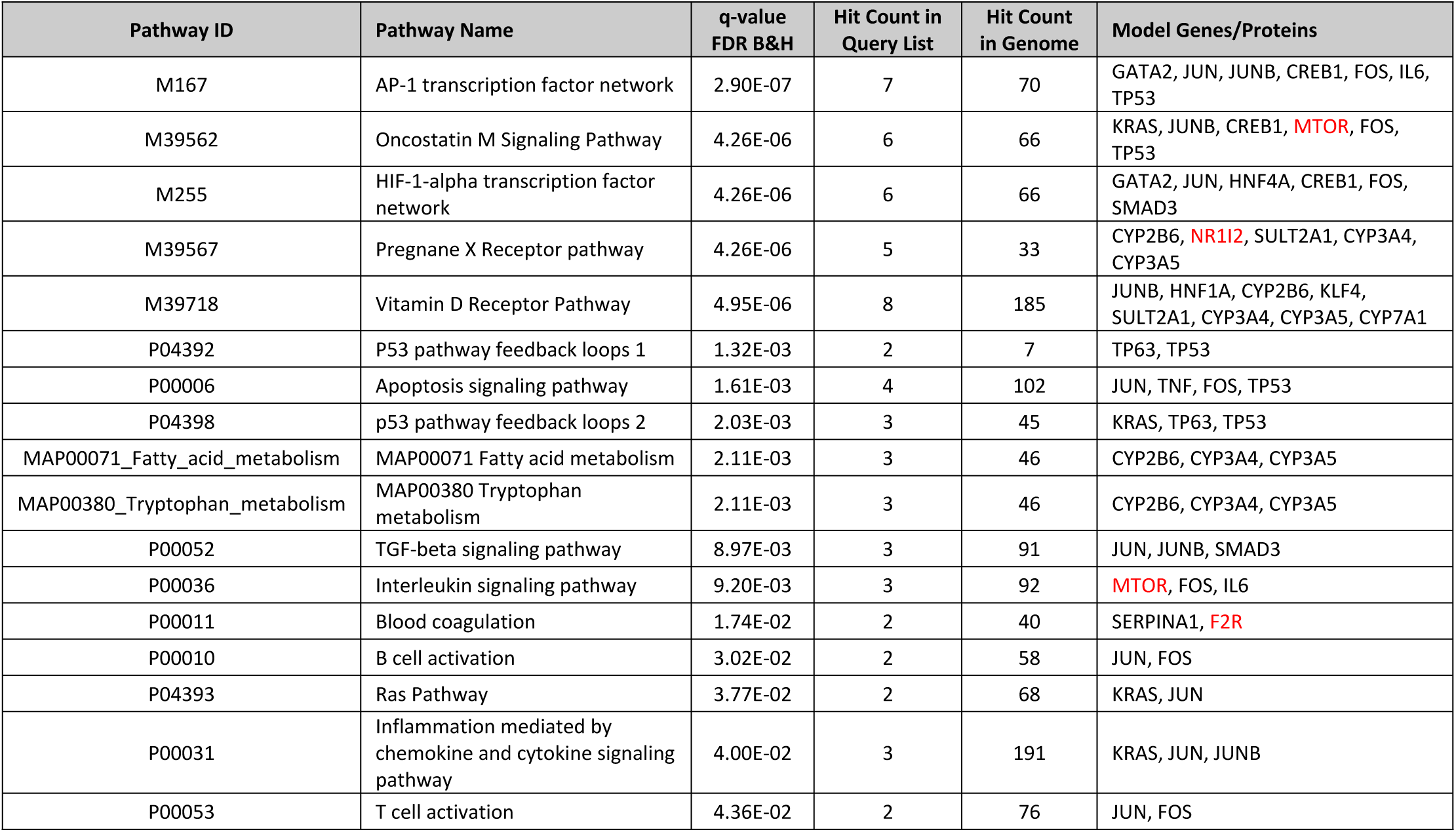
Pathways significantly enriched in the COVID-19 disease biology model. Red gene symbols indicate the COVID-19-associated genes/proteins initially identified in the literature.

| Pathway ID | Pathway Name | q-value<br>FDR B&H | Hit Count in<br>Query List | Hit Count<br>in Genome | Model Genes/Proteins |
| --- | --- | --- | --- | --- | --- |
| M167 | AP-1 transcription factor network | 2.90E-07 | 7 | 70 | GATA2, JUN, JUNB, CREB1, FOS, IL6, TP53 |
| M39562 | Oncostatin M Signaling Pathway | 4.26E-06 | 6 | 66 | KRAS, JUNB, CREB1, **MTOR**, FOS, TP53 |
| M255 | HIF-1-alpha transcription factor network | 4.26E-06 | 6 | 66 | GATA2, JUN, HNF4A, CREB1, FOS, SMAD3 |
| M39567 | Pregnane X Receptor pathway | 4.26E-06 | 5 | 33 | CYP2B6, **NR1I2**, SULT2A1, CYP3A4, CYP3A5 |
| M39718 | Vitamin D Receptor Pathway | 4.95E-06 | 8 | 185 | JUNB, HNF1A, CYP2B6, KLF4, SULT2A1, CYP3A4, CYP3A5, CYP7A1 |
| P04392 | P53 pathway feedback loops 1 | 1.32E-03 | 2 | 7 | TP63, TP53 |
| P00006 | Apoptosis signaling pathway | 1.61E-03 | 4 | 102 | JUN, TNF, FOS, TP53 |
| P04398 | p53 pathway feedback loops 2 | 2.03E-03 | 3 | 45 | KRAS, TP63, TP53 |
| MAP00071_Fatty_acid_metabolism | MAP00071 Fatty acid metabolism | 2.11E-03 | 3 | 46 | CYP2B6, CYP3A4, CYP3A5 |
| MAP00380_Tryptophan_metabolism | MAP00380 Tryptophan metabolism | 2.11E-03 | 3 | 46 | CYP2B6, CYP3A4, CYP3A5 |
| P00052 | TGF-beta signaling pathway | 8.97E-03 | 3 | 91 | JUN, JUNB, SMAD3 |
| P00036 | Interleukin signaling pathway | 9.20E-03 | 3 | 92 | **MTOR**, FOS, IL6 |
| P00011 | Blood coagulation | 1.74E-02 | 2 | 40 | SERPINA1, **F2R** |
| P00010 | B cell activation | 3.02E-02 | 2 | 58 | JUN, FOS |
| P04393 | Ras Pathway | 3.77E-02 | 2 | 68 | KRAS, JUN |
| P00031 | Inflammation mediated by chemokine and cytokine signaling pathway | 4.00E-02 | 3 | 191 | KRAS, JUN, JUNB |
| P00053 | T cell activation | 4.36E-02 | 2 | 76 | JUN, FOS |

### Disease and phenotype enrichment analysis

To validate and compare the COVID-19 and influenza A H1N1 models, disease enrichment analysis used BeFree text mining data from DisGeNET, a discovery platform containing one of the largest publicly available collections of genes and variants associated to human diseases ^28^. Both models met our validation criteria by showing significant enrichment for the modeled disease: the COVID-19 model was significantly enriched with genes/proteins associated with virus diseases (q-value FDR B&H = 2.57E-13) and coronavirus infections (q-value FDR B&H = 2.95E-03), while the H1N1 model was enriched with genes/proteins associated with virus diseases (q-value FDR B&H = 5.19E-44) and influenza A virus infection (q-value FDR B&H = 2.12E-17). The COVID-19 model was significantly enriched with genes/proteins linked to a number of additional diseases associated with COVID-19 infection (**Table 3**).

**Table 3.**
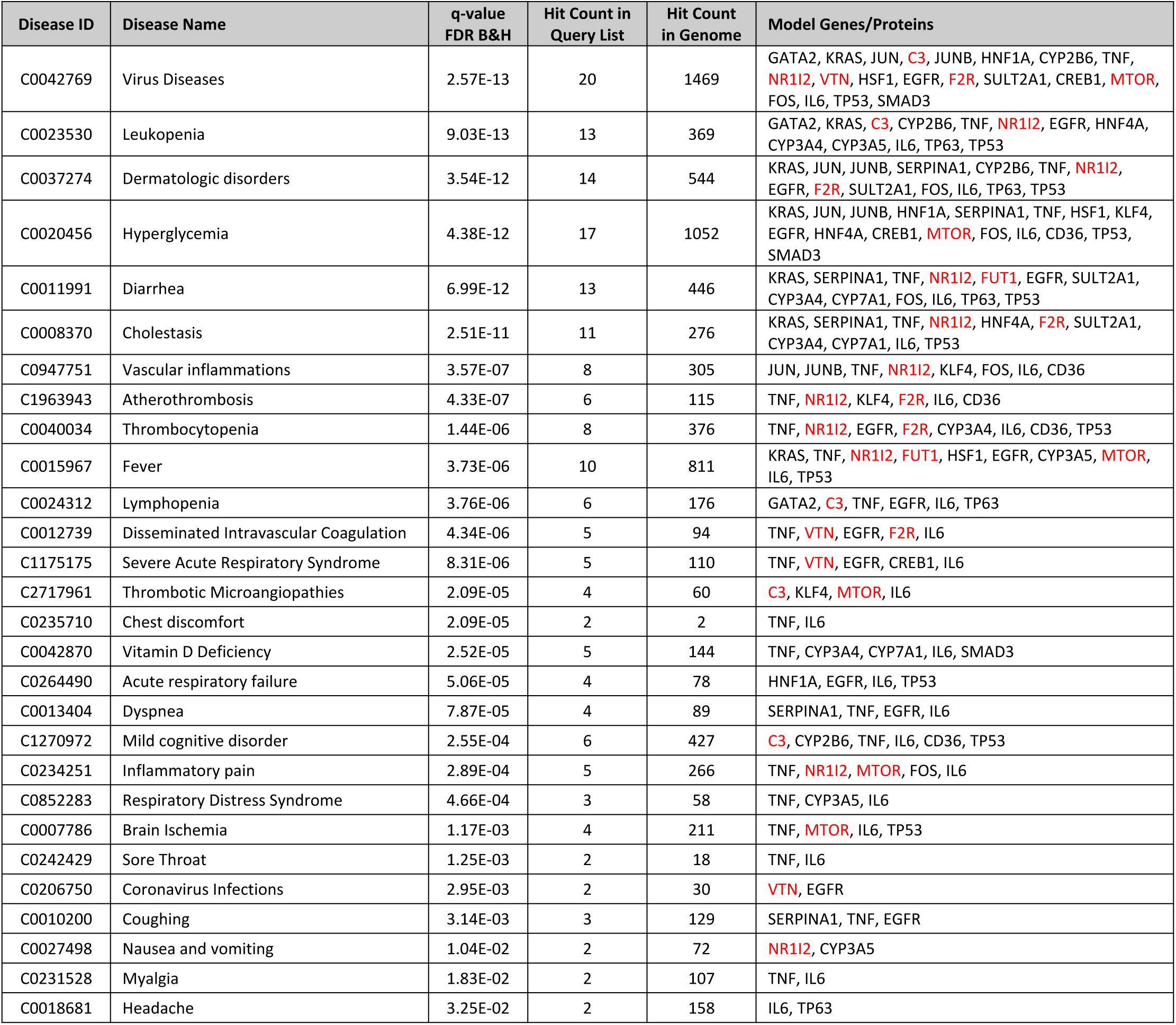
Diseases significantly enriched in the COVID-19 disease biology model. Red gene symbols indicate the COVID-19-associated genes/proteins initially identified in the literature.

| Disease ID | Disease Name | q-value<br>FDR B&H | Hit Count in<br>Query List | Hit Count<br>in Genome | Model Genes/Proteins |
| --- | --- | --- | --- | --- | --- |
| C0042769 | Virus Diseases | 2.57E-13 | 20 | 1469 | GATA2, KRAS, JUN, **C3**, JUNB, HNF1A, CYP2B6, TNF, **NR1I2**, **VTN**, HSF1, EGFR, **F2R**, SULT2A1, CREB1, **MTOR**, FOS, IL6, TP53, SMAD3 |
| C0023530 | Leukopenia | 9.03E-13 | 13 | 369 | GATA2, KRAS, **C3**, CYP2B6, TNF, **NR1I2**, EGFR, HNF4A, CYP3A4, CYP3A5, IL6, TP63, TP53 |
| C0037274 | Dermatologic disorders | 3.54E-12 | 14 | 544 | KRAS, JUN, JUNB, SERPINA1, CYP2B6, TNF, **NR1I2**, EGFR, **F2R**, SULT2A1, FOS, IL6, TP63, TP53 |
| C0020456 | Hyperglycemia | 4.38E-12 | 17 | 1052 | KRAS, JUN, JUNB, HNF1A, SERPINA1, TNF, HSF1, KLF4, EGFR, HNF4A, CREB1, **MTOR**, FOS, IL6, CD36, TP53, SMAD3 |
| C0011991 | Diarrhea | 6.99E-12 | 13 | 446 | KRAS, SERPINA1, TNF, **NR1I2**, **FUT1**, EGFR, SULT2A1, CYP3A4, CYP7A1, FOS, IL6, TP63, TP53 |
| C0008370 | Cholestasis | 2.51E-11 | 11 | 276 | KRAS, SERPINA1, TNF, **NR1I2**, HNF4A, **F2R**, SULT2A1, CYP3A4, CYP7A1, IL6, TP53 |
| C0947751 | Vascular inflammations | 3.57E-07 | 8 | 305 | JUN, JUNB, TNF, **NR1I2**, KLF4, FOS, IL6, CD36 |
| C1963943 | Atherothrombosis | 4.33E-07 | 6 | 115 | TNF, **NR1I2**, KLF4, **F2R**, IL6, CD36 |
| C0040034 | Thrombocytopenia | 1.44E-06 | 8 | 376 | TNF, **NR1I2**, EGFR, **F2R**, CYP3A4, IL6, CD36, TP53 |
| C0015967 | Fever | 3.73E-06 | 10 | 811 | KRAS, TNF, **NR1I2**, **FUT1**, HSF1, EGFR, CYP3A5, **MTOR**, IL6, TP53 |
| C0024312 | Lymphopenia | 3.76E-06 | 6 | 176 | GATA2, **C3**, TNF, EGFR, IL6, TP63 |
| C0012739 | Disseminated Intravascular Coagulation | 4.34E-06 | 5 | 94 | TNF, **VTN**, EGFR, **F2R**, IL6 |
| C1175175 | Severe Acute Respiratory Syndrome | 8.31E-06 | 5 | 110 | TNF, **VTN**, EGFR, CREB1, IL6 |
| C2717961 | Thrombotic Microangiopathies | 2.09E-05 | 4 | 60 | **C3**, KLF4, **MTOR**, IL6 |
| C0235710 | Chest discomfort | 2.09E-05 | 2 | 2 | TNF, IL6 |
| C0042870 | Vitamin D Deficiency | 2.52E-05 | 5 | 144 | TNF, CYP3A4, CYP7A1, IL6, SMAD3 |
| C0264490 | Acute respiratory failure | 5.06E-05 | 4 | 78 | HNF1A, EGFR, IL6, TP53 |
| C0013404 | Dyspnea | 7.87E-05 | 4 | 89 | SERPINA1, TNF, EGFR, IL6 |
| C1270972 | Mild cognitive disorder | 2.55E-04 | 6 | 427 | **C3**, CYP2B6, TNF, IL6, CD36, TP53 |
| C0234251 | Inflammatory pain | 2.89E-04 | 5 | 266 | TNF, **NR1I2**, **MTOR**, FOS, IL6 |
| C0852283 | Respiratory Distress Syndrome | 4.66E-04 | 3 | 58 | TNF, CYP3A5, IL6 |
| C0007786 | Brain Ischemia | 1.17E-03 | 4 | 211 | TNF, **MTOR**, IL6, TP53 |
| C0242429 | Sore Throat | 1.25E-03 | 2 | 18 | TNF, IL6 |
| C0206750 | Coronavirus Infections | 2.95E-03 | 2 | 30 | **VTN**, EGFR |
| C0010200 | Coughing | 3.14E-03 | 3 | 129 | SERPINA1, TNF, EGFR |
| C0027498 | Nausea and vomiting | 1.04E-02 | 2 | 72 | **NR1I2**, CYP3A5 |
| C0231528 | Myalgia | 1.83E-02 | 2 | 107 | TNF, IL6 |
| C0018681 | Headache | 3.25E-02 | 2 | 158 | IL6, TP63 |

The top 100 disease enrichments for each model were then mapped from DisGenNET BeFree names to DOIDs in order to compare/contrast disease classes instead of individual conditions (**Figure 2**). Indications that could be mapped to multiple high level classes were (e.g., leukemia mapped to both cancer and hematopoietic system disease). We specifically looked at acquired disease indications and their corresponding classes. The COVID-19 model was enriched with more hematopoietic system diseases, including leukopenia and neutropenia, and gastrointestinal system diseases, including diarrhea and cholestasis, than the influenza A H1N1 model. Conversely, influenza A H1N1 model was enriched with more immune system and cardiovascular system diseases than COVID-19.

**Figure 2.**
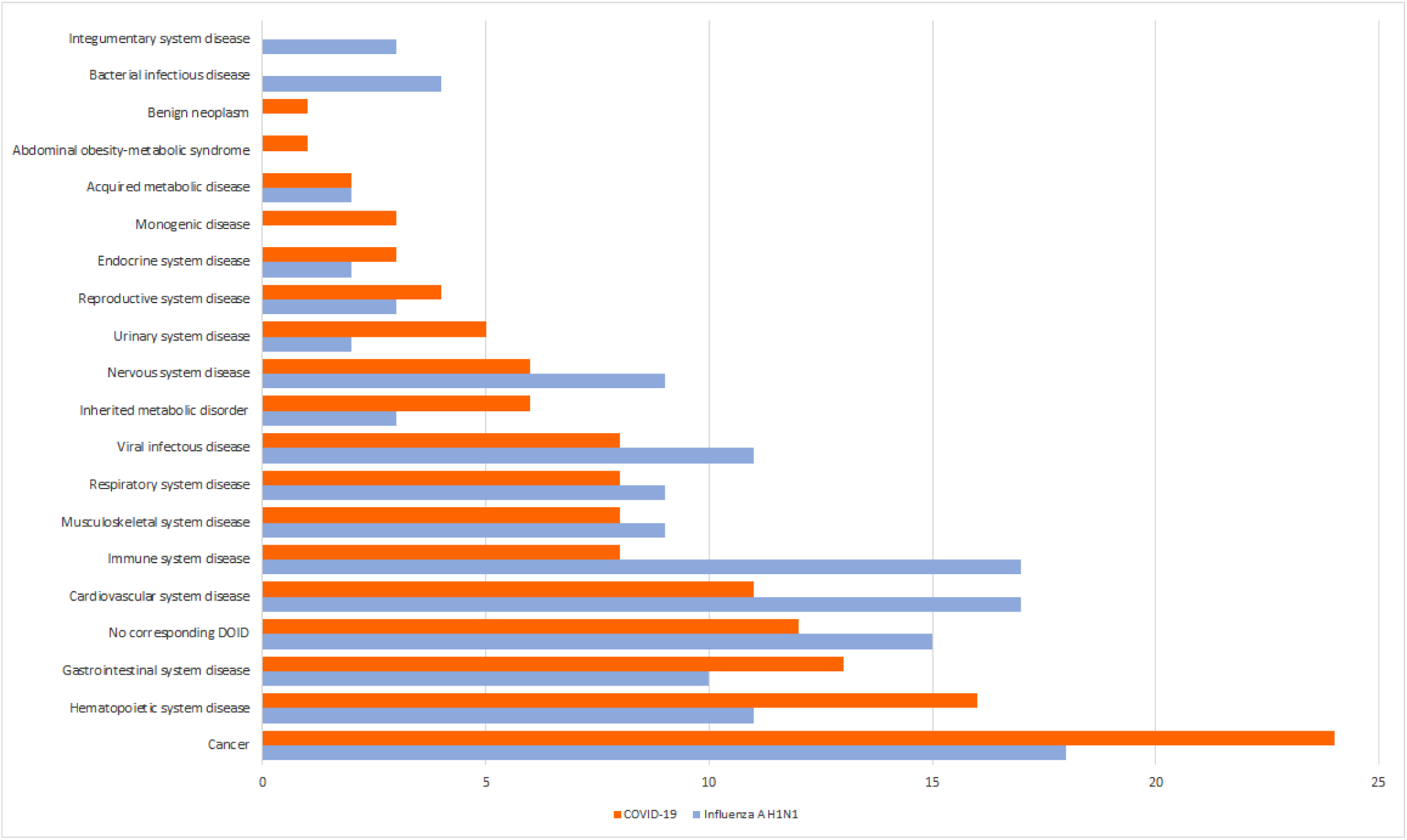
Top 100 DisGeNET BeFree COVID-19 and influenza H1N1 model disease enrichments mapped to disease ontology classes. Red: COVID-19; blue: Influenza A H1N1.

To provide additional model validation, we also performed human phenotype enrichment analysis (**Table 4**). Similar to disease enrichment, phenotype enrichments overlap published COVID-19 patient clinical presentation ^5, 29, 30^.

**Table 4.** Phenotypes significantly enriched in the COVID-19 disease biology model. Red gene symbols indicate the COVID-19-associated genes/proteins initially identified in the literature.

| Phenotype ID | Phenotype Name | q-value<br>FDR B&H | Hit Count in<br>Query List | Hit Count<br>in Genome | Model Genes/Proteins |
| --- | --- | --- | --- | --- | --- |
| HP:0000855 | Insulin resistance | 1.36E-02 | 6 | 159 | KRAS, HNF1A, HNF4A, FOS, IL6, TP53 |
| HP:0002344 | Progressive neurologic deterioration | 2.47E-02 | 4 | 67 | HNF1A, EGFR, HNF4A, IL6 |
| HP:0000818 | Abnormality of the endocrine system | 2.51E-02 | 14 | 1460 | KRAS, **C3**, HNF1A, SERPINA1, WNT4, EGFR, HNF4A, CYP3A5, FOS, IL6, TP63, CD36, TP53, SMAD3 |
| HP:0002013 | Vomiting | 2.51E-02 | 7 | 322 | KRAS, HNF1A, TNF, EGFR, HNF4A, IL6, TP53 |
| HP:0003119 | Abnormal circulating lipid concentration | 2.51E-02 | 8 | 443 | KRAS, HNF1A, SERPINA1, WNT4, HNF4A, CYP7A1, FOS, TP53 |
| HP:0010978 | Abnormality of immune system physiology | 3.84E-02 | 14 | 1554 | GATA2, KRAS, **C3**, SERPINA1, TNF, WNT4, EGFR, **MTOR**, CYP7A1, FOS, IL6, TP63, TP53, SMAD3 |
| HP:0002105 | Hemoptysis | 3.84E-02 | 5 | 163 | KRAS, SERPINA1, EGFR, TP53, SMAD3 |
| HP:0001873 | Thrombocytopenia | 3.90E-02 | 8 | 512 | GATA2, KRAS, **C3**, EGFR, **MTOR**, CD36, TP53, SMAD3 |
| HP:0100820 | Glomerulopathy | 3.90E-02 | 6 | 269 | KRAS, **C3**, SERPINA1, HNF4A, FOS, TP53 |
| HP:0012310 | Abnormal monocyte count | 3.99E-02 | 2 | 9 | GATA2, KRAS |
| HP:0001872 | Abnormal thrombocyte morphology | 3.99E-02 | 9 | 682 | GATA2, KRAS, **C3**, EGFR, **MTOR**, IL6, CD36, TP53, SMAD3 |
| HP:0002018 | Nausea | 3.99E-02 | 4 | 105 | KRAS, TNF, IL6, TP53 |
| HP:0003259 | Elevated serum creatinine | 3.99E-02 | 4 | 108 | KRAS, **C3**, HNF4A, TP53 |
| HP:0011015 | Abnormal blood glucose concentration | 3.99E-02 | 6 | 299 | KRAS, HNF1A, HNF4A, **MTOR**, FOS, TP53 |
| HP:0011873 | Abnormal platelet count | 4.03E-02 | 8 | 562 | GATA2, KRAS, **C3**, EGFR, **MTOR**, CD36, TP53, SMAD3 |
| HP:0002097 | Emphysema | 4.03E-02 | 5 | 200 | KRAS, SERPINA1, EGFR, TP53, SMAD3 |
| HP:0003117 | Abnormal circulating hormone level | 4.11E-02 | 8 | 573 | KRAS, HNF1A, WNT4, EGFR, HNF4A, FOS, TP63, TP53 |
| HP:0002014 | Diarrhea | 4.20E-02 | 7 | 439 | KRAS, HNF1A, EGFR, HNF4A, IL6, TP53, SMAD3 |

### Drug enrichment analysis

Lastly, to leverage knowledge about interactions between proteins and small molecules, we performed drug enrichment analysis, which incorporated two sources: the Comparative Toxicogenomics Database (CTD) (http://ctdbase.org/) and Stitch (http://stitch.embl.de/) (**Table 5**). We cross-referenced candidate drugs against the PubChem database to identify drug roles shown in in the table. For contextual relevance, we omitted any non-human-use compound (e.g., used as an insecticide, plastics additive, or fluorosurfactant) as well as any compound notated with a legacy record in PubChem (which indicates that the record is no longer maintained by the data contributor and may contain outdated information) or withdrawn from use due to hepatotoxicity.

**Table 5.** Drugs significantly enriched in the COVID-19 disease biology model. Drug Role antineoplastic agents highlighted in red. Model Genes/Proteins red gene symbols indicate the COVID-19-associated genes/proteins initially identified in the literature.

| Drug ID | Drug Name | PubChem ID | Drug Role | q-value<br>FDR B&H | Hit Count in<br>Query List | Hit Count<br>in Genome | Model Genes/Proteins |
| --- | --- | --- | --- | --- | --- | --- | --- |
| ctd:C113580 | U0126 (1,4-Diamino-2,3-dicyano-1,4-bis(o-aminophenylmercapto)butadiene) | 3006531 | MAP kinase inhibitor; apoptosis inducer; **antineoplastic agent**; antioxidant; osteogenesis regulator; vasoconstrictor agent | 6.48E-20 | 18 | 444 | GATA2, JUN, JUNB, CYP2B6, TNF, **NR1I2**, EGFR, HNF4A, CYP3A4, CREB1, CYP3A5, **MTOR**, CYP7A1, FOS, IL6, CD36, TP53, SMAD3 |
| ctd:D015735 | Mifepristone | 55245 | Abortifacient; contraceptive drug; synthetic oral contraceptive; hormone antagonist | 6.48E-20 | 19 | 553 | JUN, **C3**, JUNB, SERPINA1, CYP2B6, TNF, **NR1I2**, HSF1, EGFR, HNF4A, **F2R**, CYP3A4, CYP3A5, **MTOR**, CYP7A1, FOS, IL6, CD36, TP53 |
| ctd:D019821 | Simvastatin | 54454 | Anthrax lethal factor endopeptidase inhibitor; prodrug; hydroxymethylglutaryl-CoA reductase inhibitor; ferroptosis inducer | 5.54E-18 | 18 | 581 | KRAS, JUN, JUNB, HNF1A, CYP2B6, TNF, **NR1I2**, WNT4, HSF1, EGFR, CYP3A4, CREB1, CYP3A5, **MTOR**, FOS, IL6, TP53, SMAD3 |
| ctd:D001647 | Bile acids and salts | 439520 | Digestion and absorption of lipids and fat-soluble vitamins | 8.81E-17 | 11 | 79 | TNF, **NR1I2**, EGFR, HNF4A, SULT2A1, **MTOR**, CYP7A1, FOS, IL6, TP53, SMAD3 |
| ctd:C070081 | Fulvestrant | 104741 | **Antineoplastic agent**; estrogen receptor antagonist; estrogen antagonist | 4.01E-16 | 16 | 484 | JUN, **C3**, TNF, **NR1I2**, WNT4, HSF1, KLF4, EGFR, HNF4A, **F2R**, **MTOR**, FOS, IL6, CD36, TP53, SMAD3 |
| ctd:D000431 | Ethanol | 702 | Antiseptic drug; polar solvent; neurotoxin; central nervous system depressant; teratogenic agent; NMDA receptor antagonist; protein kinase C agonist; disinfectant; human metabolite; Saccharomyces cerevisiae metabolite; Escherichia coli metabolite; mouse metabolite | 4.01E-16 | 22 | 1608 | JUN, JUNB, HNF1A, CYP2B6, TNF, **NR1I2**, **VTN**, HSF1, EGFR, HNF4A, **F2R**, SULT2A1, CYP3A4, CREB1, **MTOR**, CYP7A1, FOS, IL6, CD36, TP53, INSIG1, SMAD3 |
| ctd:D003474 | Curcumin | 969516 | Metabolite; anti-inflammatory agent; **antineoplastic agent**; hepatoprotective agent; flavouring agent; biological pigment; nutraceutical; antifungal agent; dye; lipoxygenase inhibitor; ligand; radical scavenger; contraceptive drug; histone deacetylase inhibitor; immunomodulator; iron chelator; neuroprotective agent; food colouring; aldehyde reductase inhibitor; shikimate dehydrogenase inhibitor; IMP dehydrogenase inhibitor; NAD(P)H dehydrogenase (quinone) inhibitor; thioredoxin reductase inhibitor; non-specific protein-tyrosine kinase inhibitor | 1.56E-15 | 18 | 851 | GATA2, KRAS, JUN, **C3**, JUNB, TNF, **NR1I2**, HSF1, EGFR, **F2R**, CYP3A4, CREB1, **MTOR**, FOS, IL6, CD36, TP53, SMAD3 |
| CID000003285 | Ethinylestradiol | 5991 | Oral contraception; **antineoplastic agent**; anti-osteoporotic agent | 1.56E-15 | 20 | 1251 | JUN, **C3**, JUNB, HNF1A, SERPINA1, CYP2B6, TNF, **NR1I2**, WNT4, HSF1, EGFR, SULT2A1, CYP3A4, CREB1, CYP3A5, CYP7A1, FOS, IL6, CD36, TP53 |
| ctd:D003687 | Dehydroepiandrosterone | 5881 | Androgen; human metabolite; mouse metabolite | 1.82E-14 | 11 | 141 | KRAS, JUN, TNF, **NR1I2**, EGFR, SULT2A1, CYP3A4, CYP3A5, FOS, IL6, TP53 |
| ctd:C103303 | Alitretinoin | 449171 | **Antineoplastic agent**; retinoid X receptor agonist; metabolite; keratolytic drug | 2.67E-14 | 13 | 300 | JUN, TNF, **NR1I2**, EGFR, HNF4A, SULT2A1, CYP3A4, CREB1, **MTOR**, CYP7A1, IL6, CD36, SMAD3 |
| ctd:C432165 | Pyrazolanthrone | 8515 | c-Jun N-terminal kinase inhibitor | 6.33E-14 | 13 | 322 | GATA2, JUN, TNF, HSF1, EGFR, HNF4A, CYP3A4, CREB1, FOS, IL6, CD36, TP53, SMAD3 |
| ctd:D008095 | Lithocholic acid | 9903 | Human metabolite; mouse metabolite | 9.19E-14 | 12 | 240 | JUN, JUNB, HNF1A, TNF, **NR1I2**, HNF4A, SULT2A1, CYP3A4, CREB1, CYP7A1, IL6, SMAD3 |
| ctd:D003022 | Clotrimazole | 2812 | Antifungal agent; environmental contaminant; xenobiotic | 9.35E-14 | 9 | 66 | CYP2B6, **NR1I2**, HNF4A, SULT2A1, CYP3A4, CYP3A5, CYP7A1, FOS, TP53 |
| ctd:D019438 | Ritonavir | 392622 | Antiviral drug; HIV protease inhibitor; environmental contaminant; xenobiotic | 9.35E-14 | 10 | 109 | JUN, CYP2B6, TNF, **NR1I2**, CYP3A4, CYP3A5, CYP7A1, FOS, IL6, TP53 |
| ctd:D005978 | Glutathione | 124886 | Skin lightening agent; human metabolite; Escherichia coli metabolite; mouse metabolite; antioxidant; cofactor | 1.00E-13 | 13 | 339 | JUN, JUNB, CYP2B6, TNF, KLF4, EGFR, HNF4A, CYP3A4, CYP3A5, FOS, IL6, CD36, TP53 |
| ctd:D020123 | Sirolimus | 5284616 | Immunosuppressive agent; **antineoplastic agent**; antibacterial drug; mTOR inhibitor; bacterial metabolite; anticoronaviral agent; geroprotector | 1.05E-13 | 14 | 456 | JUN, **C3**, SERPINA1, CYP2B6, TNF, EGFR, CYP3A4, CREB1, **MTOR**, IL6, TP63, CD36, TP53, INSIG1 |
| ctd:C043561 | Chrysin | 5281607 | Anti-inflammatory agent; **antineoplastic agent**; antioxidant; hepatoprotective agent; myosin-light-chain kinase inhibitor; plant metabolite | 1.41E-13 | 10 | 115 | JUN, CYP2B6, TNF, EGFR, CYP3A4, FOS, IL6, TP63, TP53, SMAD3 |
| ctd:D000111 | Acetylcysteine | 12035 | Mucolytic; hepatoprotective agent; antioxidant | 1.69E-13 | 16 | 781 | KRAS, JUN, **C3**, JUNB, TNF, HSF1, EGFR, CYP3A4, CREB1, **MTOR**, FOS, IL6, CD36, TP53, INSIG1, SMAD3 |
| CID000000784 | H2O2 (Hydrogen Peroxide) | 784 | Antiseptic drug | 1.79E-13 | 18 | 1183 | JUN, JUNB, SERPINA1, TNF, HSF1, KLF4, EGFR, **F2R**, CYP3A4, CREB1, CYP3A5, **MTOR**, FOS, IL6, TP63, CD36, TP53, SMAD3 |
| ctd:D002110 | Caffeine | 2519 | Central nervous system stimulant; phosphoric diester hydrolase inhibitor; adenosine receptor antagonist; non-specific serine/threonine protein kinase inhibitor; ryanodine receptor agonist; fungal metabolite; adenosine A2A receptor antagonist; psychotropic drug; diuretic; food additive; adjuvant; plant metabolite; environmental contaminant; xenobiotic; human blood serum metabolite; mouse metabolite; mutagen | 4.04E-13 | 14 | 515 | JUN, SERPINA1, TNF, **VTN**, WNT4, KLF4, CYP3A4, CREB1, **MTOR**, FOS, IL6, TP53, INSIG1, SMAD3 |
| ctd:D047311 | Luteolin | 5280445 | Fatty acid synthase inhibitor; **antineoplastic agent**; vascular endothelial growth factor receptor antagonist; plant metabolite; nephroprotective agent; angiogenesis inhibitor; c-Jun N-terminal kinase inhibitor; anti-inflammatory agent; apoptosis inducer; radical scavenger; immunomodulator | 4.07E-13 | 10 | 132 | JUN, JUNB, TNF, EGFR, CYP3A4, CYP3A5, FOS, IL6, TP63, TP53 |
| CID000003143 | Docetaxel trihydrate | 148123 | **Antineoplastic agent** | 4.53E-13 | 11 | 202 | JUN, HNF1A, CYP2B6, **NR1I2**, EGFR, HNF4A, CYP3A4, CYP3A5, **MTOR**, IL6, TP53 |
| ctd:C093973 | 2-(2-amino-3-methoxyphenyl)-4H-1-benzopyran-4-one | 4713 | MAP kinase inhibitor | 5.47E-13 | 13 | 402 | GATA2, KRAS, JUN, JUNB, TNF, HSF1, EGFR, CREB1, CYP7A1, FOS, IL6, CD36, TP53 |
| ctd:D000661 | Amphetamine | 3007 | Psychotropic drug | 1.11E-12 | 10 | 148 | JUN, JUNB, CYP2B6, TNF, **NR1I2**, CYP3A4, CREB1, FOS, IL6, TP53 |
| ctd:D017239 | Paclitaxel | 36314 | Microtubule-stabilising agent; metabolite; human metabolite; **antineoplastic agent** | 1.11E-12 | 15 | 720 | JUN, **C3**, TNF, **NR1I2**, WNT4, EGFR, SULT2A1, CYP3A4, **MTOR**, CYP7A1, IL6, CD36, TP53, INSIG1, SMAD3 |
| ctd:D020148 | Butyric Acid | 264 | Mycoplasma genitalium metabolite; human urinary metabolite | 2.49E-12 | 10 | 161 | GATA2, JUN, JUNB, HNF1A, TNF, **MTOR**, FOS, IL6, TP53, INSIG1 |
| ctd:D013792 | Thalidomide | 5426 | Immunomodulator; **antineoplastic agent**; anti-inflammatory agent; angiogenesis inhibitor | 4.25E-12 | 13 | 479 | JUN, **C3**, CYP2B6, TNF, **NR1I2**, **VTN**, HSF1, **F2R**, CYP3A4, CYP3A5, FOS, IL6, TP53 |
| ctd:D012293 | Rifampin | 135398735 | RNA polymerase inhibitor; DNA synthesis inhibitor; antitubercular agent; leprostatic drug; Escherichia coli metabolite; protein synthesis inhibitor; neuroprotective agent; angiogenesis inhibitor; pregnane X receptor agonist; **antineoplastic agent**; antiamoebic agent | 4.30E-12 | 12 | 356 | JUN, CYP2B6, TNF, **NR1I2**, **FUT1**, HNF4A, SULT2A1, CYP3A4, CYP3A5, CYP7A1, IL6, CD36 |
| ctd:D003915 | Dextromethorphan | 5360696 | NMDA receptor antagonist; neurotoxin; xenobiotic; environmental contaminant; antitussive; prodrug; oneirogen | 6.50E-12 | 7 | 34 | JUN, CYP2B6, TNF, CYP3A4, CYP3A5, FOS, IL6 |
| ctd:C037032 | Galangin | 5281616 | Antimicrobial agent; triacylglycerol lipase inhibitor; plant metabolite | 8.42E-12 | 8 | 68 | CYP2B6, TNF, **NR1I2**, EGFR, CYP3A4, **MTOR**, IL6, SMAD3 |

From the top thirty drug candidates, twelve have a role as an antineoplastic agent. We queried each against the U.S. Food and Drug Administration (FDA) Drugs@FDA database and seven are approved for human use: fulvestrant and ethylnylestradiol target the estrogen receptor; paclitaxel stabilizes tubulin, inhibits the disassembly of microtubules, and blocks mitosis; alitretinoin is a retinoid that binds to and activates nuclear retinoic acid receptors (RAR) and retinoid X receptors (RXR); sirolimus is a macrocyclic lactone that binds to the immunophilin FK Binding Protein-12 (FKBP-12) to generate an immunosuppressive complex that binds to and inhibits the activation of the mammalian Target Of Rapamycin (mTOR), one of the 6 seed genes/proteins identified in the literature and used to build the COVID-19 disease biology model; rifampin inhibits DNA-dependent RNA polymerase in susceptible bacteria; and thalidomide, an immunomodulatory agent that inhibits both the production of tumor necrosis factor alpha (TNF-alpha) in stimulated peripheral monocytes and the activities of interleukins and interferons, inhibits polymorphonuclear chemotaxis and monocyte phagocytosis, and inhibits pro-angiogenic factors such as vascular endothelial growth factor (VEGF) and basic fibroblast growth factor (bFGF), thereby inhibiting angiogenesis. Of the FDA-approved antineoplastic agents, three are planned or in one or more COVID-19 clinical trials as the primary drug being investigated (number of trials in parentheses): alitretinoin (1), sirolimus (7) and thalidomide (7). The current status of these trials is shown in **Table 6**.

**Table 6.** COVID-19 clinical trial status with model-derived FDA-approved antineoplastic agents.

| Drug Name | Clinical trials |  |  |  |  |  | Total Number, All Statuses |
| --- | --- | --- | --- | --- | --- | --- | --- |
|  | Open | Planned | Completed | Terminated | Closed | Temporarily Closed |  |
| Alitretinoin | 0 | 1 | 0 | 0 | 0 | 0 | 1 |
| Sirolimus | 3 | 1 | 0 | 3 | 0 | 0 | 7 |
| Thalidomide | 1 | 3 | 2 | 0 | 1 | 0 | 7 |

There are also clinical trials with non-antineoplastic agents as the primary drug being investigated for the treatment of COVID-19 (number of trials in parentheses): simvastatin (3), ethanol (5), curcumin (17), clotrimazole (18), ritonavir (272), glutathione (6), N-acetylcysteine (37), and hydrogen peroxide (20). A subset of these agents and trials are being investigated specifically for moderate to severe or critical COVID-19 patients (number of trials in parentheses): simvastatin (3), ethanol (2), curcumin (5), clotrimazole (8), ritonavir (125), glutathione (7), sirolimus (6), N-acetylcysteine (14) and thalidomide (7); these are shown in **Table 7**.

**Table 7.** Clinical trial status with model-derived agents in moderate to severe or critical COVID-19 patients.

| Drug Name | Clinical trials |  |  |  |  |  | Total Number, All Statuses |
| --- | --- | --- | --- | --- | --- | --- | --- |
|  | Open | Planned | Completed | Terminated | Closed | Temporarily Closed |  |
| Simvastatin | 1 | 0 | 0 | 1 | 1 | 0 | 3 |
| Ethanol | 0 | 1 | 1 | 0 | 0 | 0 | 2 |
| Curcumin | 3 | 0 | 1 | 0 | 1 | 0 | 5 |
| Clotrimazole | 3 | 2 | 2 | 1 | 0 | 0 | 8 |
| Ritonavir | 40 | 14 | 50 | 7 | 12 | 2 | 125 |
| Glutathione | 3 | 1 | 1 | 1 | 1 | 0 | 7 |
| Sirolimus | 3 | 1 | 0 | 2 | 0 | 0 | 6 |
| N-Acetylcysteine | 4 | 2 | 5 | 2 | 1 | 0 | 14 |
| Thalidomide | 1 | 3 | 2 | 0 | 1 | 0 | 7 |

## Discussion

We report here the development and analysis of a COVID-19 disease biology model. The model is based on genes/proteins identified in the research literature together with known molecular interactions in epithelial cells (**Figure 1**). We followed a systematic and reproducible strategy to construct the model, and did not manually add genes/proteins, nor include PPIs not present in the knowledgebase. For example, ACE2 is missing from the final model even though it was in the initial list of 14 COVID-19- associated genes/proteins identified in the research literature. This is due to a limitation in the IPA knowledgebase; there are no direct physical interactions in epithelial cells up- or down-stream of ACE2 in the knowledgebase. Thus, since it does not connect to any other nodes in our model, it was not included.

### Host biological response induced by SARS-CoV-2

Biological process and pathway enrichment analyses provide context to better understand disease biology (**Tables 1, 2**). At a high level, these analyses reflect viral infection by way of xenobiotic metabolism, host cellular damage (i.e. response to wounding), defense response and cytokine production, and a change in extracellular-signal-regulated kinase (ERK) signaling, the last step of the Ras/Raf/Mitogen-activated protein kinase/ERK kinase (MEK)/ERK signal transduction pathway. Upon various extracellular stimuli, this regulatory cascade results in ERK-1 and ERK-2 activation, which phosphorylates numerous downstream substrates and leads to the expression of multiple genes essential for diverse cellular functions, such as cell proliferation, differentiation, survival or apoptosis ^31, 32^. Many of these functions were captured by our model, including apoptosis signaling and negative regulation of apoptotic process, the Ras pathway, leukocyte differentiation, response to oxidative stress, and P53 pathway feedback loops (**Tables 1, 2**). Notably, our model captures some of the hematological and neurological manifestations of COVID-19 infection, including blood microparticle formation, coagulation, and cognition.

Transforming growth factor beta (TGF-beta) activates ERK signaling ^33^. The TGF-beta signaling pathway is involved in many cellular processes including cell survival, apoptosis and immunity. The TGF-beta signaling pathway was over-represented in our COVID-19 disease biology model (**Table 2**). TGF-beta has been implicated as a key cytokine regulating the ongoing immune activation in severe COVID-19 ^34^ and limits antiviral functions of NK cells ^35^.

ERK activation leads to elevated AP-1 activity via c-fos induction ^36^. In our COVID-19 disease biology model, the AP-1 transcription factor network is significantly over-represented (**Table 1**). Zhu et al. recently compared drug screen activity profiles with the drug activity profile in a cytopathic effect assay of SARS-CoV-2 and found that the autophagy and AP-1 signaling pathway activity profiles are significantly correlated with the anti-SARS-CoV-2 activity profile ^37^. AP-1 refers to a group of transcription factors that regulates gene expression in response to a variety of stimuli, including cytokines, growth factors, stress, and bacterial and viral infections ^38^. The AP-1 family consists of hetero- and homodimers of bZIP (basic region leucine zipper) proteins, mainly of Jun-Jun, Jun-Fos, or Jun-ATF. AP-1 regulation of cellular processes, including cell proliferation, death, survival and differentiation, critically depends on the relative abundance of AP-1 subunits, the composition of AP-1 dimers, the quality of stimulus, the cell type, and co-factor assembly. Further, concordant with identification of autophagy by Zhu et al., we identified the biological process of membrane invagination significantly enriched in the COVID-19 disease biology model (**Table 2**). The mechanism and molecules involved in membrane invagination during virus endocytosis are unknown. Autophagy has been reported in other coronavirus infections ^39–41^, and two drugs in our list of drug candidates that could potentially be repurposed to treat COVID-19 infection – U0126 (1,4-Diamino-2,3-dicyano-1,4-bis(o-aminophenylmercapto)butadiene) and 2-(2-amino-3-methoxyphenyl)-4H-1-benzopyran-4-one – inhibit autophagy.

The oncostatin M (OSM) signaling pathway was over-represented in our COVID-19 disease biology model (**Table 2**). OSM, a multifunctional cytokine that belongs to the interleukin-6 (IL6) subfamily ^42^, was found to be one of the most potent proinflammatory inducers in lung-derived epithelial cells ^43^, and in liver epithelial cells, OSM enhances the antiviral effects of type I interferon and activates immunostimulatory functions ^44^. Elevated levels of OSM have been found in the serum of COVID-19 patients in intensive care units that correlated with disease severity ^45^. In macrophages, a functional AP-1 site is required for OSM activation by thrombin ^46^. There may be similar involvement of both AP-1 and OSM in activation of the inflammatory response to SARS-CoV-2 infection and/or cytokine storm reactions in patients with severe disease. Along with OSM signaling, inflammation mediated by chemokine and cytokine signaling pathway was also over-represented in our COVID-19 disease biology model (**Table 2**).

In our biological process enrichment analysis, cellular response to oxidative stress, inflammatory response, regulation of immune system process, and positive regulation of receptor signaling pathway via JAK-STAT (Janus kinase/signal transducers and activators of transcription) were all significantly over-represented in our model (**Table 1**). The JAK/STAT signaling pathway mediates cellular responses to cytokines and growth factors, and is involved in orchestrating hematopoiesis, inducing inflammation, and controlling the immune response ^47^. The JAK/STAT pathway is implicated in the SARS-CoV-2-induced cytokine storm ^48^, and several studies have suggested targeting the pathway to control cytokine release syndrome in COVID-19 ^49–53^.

Along with JAK/STAT signaling, the inflammation interleukin signaling pathway was also over-represented in our COVID-19 disease biology model (**Table 2**). This pathway contains JAK/STAT, as well as the Ras/Raf/MEK/ERK cascade, which couples signals from cell surface receptors to transcription factors to regulate gene expression. The Porcine deltacoronavirus (PDCoV), a newly emerged swine coronavirus that causes acute enteritis in neonatal piglets, was found to activate the Raf/MEK/ERK pathway to promote its replication ^54^. Our drug enrichment analysis identified eight drug candidates that play a role as an antioxidant, anti-inflammatory, or immunosuppressive agent – U0126 (1,4-Diamino-2,3-dicyano-1,4-bis(o-aminophenylmercapto)butadiene), curcumin, glutathione, sirolimus, chrysin, acetylcysteine, luteolin, and thalidomide – and could potentially be repurposed to treat COVID-19 infection.

In our COVID-19 disease biology model, over-representation of the HIF-1-alpha transcription factor network included HNF4A. The HNF4A protein regulates genes that are important for development and function of beta cells in the pancreas, which produce and secrete insulin. Dysfunction of HNF4A has been associated with diabetes mellitus ^55–58^. HNF4A expression is also detected in epithelia of the pancreas, kidneys, stomach, and intestine, where it exerts functional roles in regulating epithelial junctions and cell proliferation. HNF4A has been found to be a transcriptional sensor of inflammation in the liver and gastrointestinal tract ^59^. It also plays a key role in the regulation of angiotensinogen metabolism ^60^ and has been predicted to regulate intestinal ACE2 expression ^61^.

Vitamin D status is associated with risks of influenza and respiratory tract infections, having direct antiviral effects primarily against enveloped viruses, including COVID-19 ^62^. Gut epithelial vitamin D receptor (VDR) is important in protecting mucosal barrier integrity and regulating the gut innate immunity ^63^. In our COVID-19 disease biology model, the VDR pathway was over-represented. A recent systematic review of vitamin D and COVID-19 found that COVID-19 infection, prognosis, and mortality were correlated with vitamin D status ^64^.

The Pregnane X receptor (PXR) pathway, also known as the steroid and xenobiotic sensing nuclear receptor (SXR) or nuclear receptor subfamily 1, group I, member 2 (NR1I2), is closely related to VDR and was over-represented in our COVID-19 disease biology model. PXR is a ligand-inducible master transcription factor regulating xenobiotic- and drug-inducible expression of key genes that encode members of the phase I and phase II metabolic enzymes and drug transporters. PXR is expressed mainly in cells of the gastrointestinal tract and liver ^65^, and has around 60% homology with VDR in their DNA-binding domains. Thus, PXR can bind to vitamin D-responsive elements in DNA and, as a transcription factor, affect the expression of genes whose expression is normally regulated by vitamin D. PXR activation has been found to suppress the activity of NF-kB, the key regulator of inflammation and immune response ^66–69^. Ligand-activated PXR has been found to interfere with HNF4A signaling ^70, 71^, establishing a link between three pathways identified in our pathway enrichment analysis: the HIF-1-alpha transcription factor network, VDR pathway, and PXR pathway. Notably, VDR, PXR, and CAR are all members of the NR1I subfamily of nuclear hormone receptors. NR1I subfamily members all share the ability to bind potentially toxic endogenous compounds with low affinity and play major roles in metabolizing xenobiotics and endogenous compounds ^72^.

### Drug enrichment analysis

The drugs suggested by the model outputs include seven compounds, three of which are FDA-approved antineoplastic agents (**Table 6**), that have been trialed, are being investigated in current trials, or are planned to be investigated in trials that have not yet started. Including other candidate drugs and agents being evaluated for use in moderate to severe or critical COVID-19 patients (**Table 7**), nearly 40% of the top 30 drugs identified by our model have been identified by others as being suitable for trialing in humans for SARS-CoV-2 infection.

The semisynthetic antibiotic rifampin is a DNA-dependent RNA polymerase inhibitor and front-line treatment for tuberculosis. Rifampin also binds PXR and is an established PXR agonist commonly used as a prototypical PXR-activating compound ^73^; the PXR pathway was significantly enriched in our pathway analysis (**Table 2**). In addition, rifampin displays antineoplastic activity *in vitro* ^74, 75^. Rifampin has been identified as a SARS-CoV-2 RNA-dependent RNA polymerase inhibitor using a cell-based SARS-CoV-2 RNA-dependent RNA polymerase activity assay system ^76^. A number of *in silico* screening studies of potent inhibitors against SARS-CoV-2 have also identified rifampin ^77–81^. Rifampicin was also found to have good binding affinity with three cytokines involved in cytokine storm (TNF-alpha, IL-6, and IL-1-beta) ^80^, suggesting that it may have a poly-pharmacological effect in COVID-19 patients.

Another antineoplastic drug candidate, U0126, inhibits influenza A virus propagation by inducing nuclear retention of the viral ribonucleoprotein complexes (RNPs), impaired function of the nuclear export protein (NEP/NS2), and concomitant inhibition of virus production ^82^. U0126 also interferes with the spread of Borna disease virus (BDV) to neighboring cells ^83^. Swine acute diarrhea syndrome coronavirus (SADS-CoV) is a newly discovered enteric coronavirus and treatment of cells with U0126 significantly inhibited SADS-CoV infection ^84^. U0126 is not currently approved by the FDA for animal or human use.

Various therapeutic effects of curcumin have been reported, including potential chemotherapeutic, antioxidant, antiviral, antibacterial, and anti-inflammatory properties ^85^. Nanoencapsulated curcumin treatment in patients with mild and severe disease resulted in a significant reduction in clinical manifestations of COVID-19 (e.g., fever, cough, and dyspnea) and reduced patient mortality rate ^86, 87^. The effectiveness of curcumin-containing nanomicelles as a therapeutic agent for the treatment of COVID-19 is currently being pursued in 17 clinical trials, five of which are for moderate to severe or critical COVID-19 (**Table 7**).

Candidate non-antineoplastic agents enriched in the model may also be useful for the treatment of COVID-19. For example, simvastatin, widely used as a hydroxymethylglutaryl (HMG)-CoA reductase inhibitor to lower blood cholesterol, was associated with a lower risk of developing severe COVID-19 and a faster time to recovery among hospitalized patients without severe disease if used during the 30 days prior to hospital admission ^88^. In a retrospective observational study of hospitalized COVID-19 patients taking simvastatin completed in 2020 (ClinicalTrials.gov identifier NCT04407273) there was a significantly lower mortality rate in patients on statin therapy than the non-statin group, and the mortality rate was even lower in patients who maintained their statin treatments during hospitalization ^89^. Three trials of simvastatin for COVID-19 treatment, all for moderate to severe or critical COVID-19, are currently being pursued (**Table 7**).

Acetylcysteine, also known as N-acetylcysteine (NAC), is a precursor of the antioxidant glutathione (another drug candidate in **Table 5**) and has been in clinical use for more than 50 years as a mucolytic drug. NAC treatment in patients with influenza was found to significantly decrease the frequency of influenza-like episodes, as well as the severity and duration of most symptoms ^90^. Subsequent research found that NAC inhibited human influenza virus (H5N1, Vietnam/VN1203 strain) replication in human lung epithelial cells in a dose-dependent manner and reduced the production of pro-inflammatory cytokines, thus reducing chemotactic migration of monocytes ^91^. Thirty-seven clinical trials of NAC, fourteen of which are for moderate to severe or critical COVID-19 (**Table 7**), are currently being pursued.

### Disease biology model limitations

There are several limitations to our COVID-19 disease biology model. The discovery of relevant molecular targets and protein interactions is dependent on existing knowledge; the model cannot make use of existing biological knowledge if it isn’t cataloged in the knowledgebase. On that note, scientific rigor and data quality of published research can have a direct impact on the model; we rely on the ensemble of molecules and interactions to outweigh any potentially poor quality data inputs. As COVID-19 can impact multiple organs/tissue types, our model may fail to identify important PPIs because they cannot be recapitulated in epithelial cells. Lastly, our model is a static representation of interactions, neglecting the temporal and spatial organization of protein dynamics.

## Conclusion

In conclusion, our study not only presents a systematic examination of host targets and cellular networks perturbed in COVID-19 patients, revealing insights into the mechanisms by which SARS-CoV-2 can trigger a cytokine storm, but also provides a list of candidate drugs, representing a powerful resource in the pursuit of therapeutic interventions.

## List of Abbreviations Abbreviations

ARDS: acute respiratory distress syndrome
bFGF: basic fibroblast growth factor
BDV: Borna disease virus
COVID-19: coronavirus disease 2019
CTD: Comparative Toxicogenomics Database
DAG: directed acyclic graph
DOID: Disease Ontology ID
ERK: extracellular-signal-regulated kinase
EUA: emergency use authorization
FDA: Food and Drug Administration
FKBP-12: FK Binding Protein-12
GenMAPP: Gene Map Annotator and Pathway Profiler
GO: Gene Ontology
GO:BP: Gene Ontology biological process
HMG: hydroxymethylglutaryl
HPO: Human phenotype ontology
IL-6: interleukin 6
IL-6R: interleukin 6 receptor
JAK: Janus kinase
JAK-STAT: Janus kinase/signal transducers and activators of transcription
MEK: mitogen-activated protein kinase/ERK kinase
mTOR: mammalian Target Of Rapamycin
NAC: N-acetylcysteine
NEP/NS2: nuclear export protein
NIH: National Institutes of Health
NR1I2: nuclear receptor subfamily 1, group I, member 2
OSM: oncostatin M
PDCoV: porcine deltacoronavirus
PPI: protein-protein interaction
PXR: pregnane X receptor
RAR: retinoic acid receptor
RNP: ribonucleoprotein complex
RXR: retinoid X receptor
SADS-CoV: Swine acute diarrhea syndrome coronavirus
SARS-CoV-2: severe acute respiratory syndrome coronavirus 2
SXR: steroid and xenobiotic sensing nuclear receptor
TGF-beta: transforming growth factor beta
TNF-alpha: tumor necrosis factor alpha
VDR: vitamin D receptor
VEGF: vascular endothelial growth factor

## Acknowledgements

No grants or financial support were used for this study.

## Conflict of interest

All authors are employed by Laboratory Corporation of America Holdings (Labcorp), some of whom own Labcorp stock.

## Notes

### Competing Interest Statement

The authors have declared no competing interest.

